# Three-photon light-sheet fluorescence microscopy

**DOI:** 10.1101/323790

**Authors:** Adriá Escobet-Montalbán, Federico M. Gasparoli, Jonathan Nylk, Pengfei Liu, Zhengyi Yang, Kishan Dholakia

## Abstract

We present the first demonstration of three-photon excitation light-sheet fluorescence microscopy. Light-sheet fluorescence microscopy in single- and two-photon modes has emerged as a powerful wide-field, low photo-damage technique for fast volumetric imaging of biological samples. We extend this imaging modality to the three-photon regime enhancing its penetration depth. Our present study uses a standard conventional femtosecond pulsed laser at 1000 nm wavelength for the imaging of 450 µm diameter cellular spheroids. In addition, we show, experimentally and through numerical simulations, the potential advantages in three-photon light-sheet microscopy of using propagation-invariant Bessel beams in preference to Gaussian beams.

---

Over the last two decades, the field of fluorescence microscopy has witnessed remarkable developments including super-resolution and fast volumetric imaging among many other innovations. However, a key remaining challenge is to perform imaging in situations where the scattering of light limits the penetration and performance of optical microscopy. This is crucial for imaging minute details of live biological samples at depth, without compromising their viability.

To increase depth penetration, multiphoton microscopy has come to the fore particularly in the form of two-photon (2P) excitation microscopy which has become the approach of choice for *in vivo* imaging [1, 2]. Recently, three-photon (3P) excitation microscopy with either point scanning [3] or with temporal focusing [4] has been employed to excite fluorophores with close to diffraction limited resolution into biological tissue for a greater penetration depth. Compared to standard single-photon (1P) or 2P excitation, 3P has several benefits: the use of longer wavelengths reduces the effects of light scattering, increasing the penetration depth of the illumination beam into the sample [3, 5]. Moreover, the nonlinear nature of the process confines the excitation to a smaller volume, reducing out-of-focus light as well as minimizing photo-bleaching on the biological sample [3, 6].

In parallel, the geometry used in light-sheet fluorescence microscopy (LSFM) has revolutionized the field of imaging by using a thin sheet of light to optically section samples which are typically transparent. In this technique, fluorescent light emitted by the sample is collected by a detection imaging system that is perpendicular to the illuminated plane. This particular configuration results in improved contrast and high axial resolution with very short acquisition times because it avoids scanning a focused beam across the field-of-view (FOV) [7]. In addition, as only the plane of interest is illuminated during a single exposure, photo-toxicity is vastly reduced. This makes LSFM very attractive for long term live imaging of biomedical samples [8, 9]. At the same time, the FOV can be increased in LFSM notably by using propagation invariant light fields such as Bessel and Airy beams [10, 11].

In this letter, we present the first demonstration of LSFM using 3P excitation. Our goal in the present work is to provide an approach to achieve greater imaging depths for biomedical imaging and explore advantages over the 2P counterpart in this particular imaging mode. The majority of research in the field of 3P microscopy has been performed using ultrashort pulsed lasers in imaging windows centered around wavelengths of 1300 nm and 1700 nm with pulse duration and repetition rate below 70 fs and 1.25 MHz [3–5, 12–14], respectively. However, such lasers are not available in most biomedical laboratories and they are limited to the imaging of green and red fluorescent samples, respectively. In this study we use a conventional Ti:Sapphire ultrashort pulsed laser, normally used for 2P microscopy, to generate 3P excitation of fluorophores with 1P absorption peaks in the violet and UV region of the spectrum (*λ* < 400 nm), including a PUREBLU™ Hoechst 33342 dye (Bio-Rad) and blue fluorescing polymer microspheres (B0100, 1 µm, Duke Scientific). The laser used in our investigation is a Coherent Chameleon Ultra II with tunable central wavelength between 680 nm and 1080 nm, 140 fs pulse duration and 80 MHz repetition rate. The use of long pulse duration and high repetition rate results in a reduced pulse energy that makes 3P excitation less efficient compared to other laser sources reported in the literature. Consequently, high average power has to be delivered to the sample which may result in increased photo-damage. In this investigation we do not focus on the optimization of 3P excitation efficiency.

3P fluorescence scales with the third power of the illumination intensity [15]. This was confirmed by measuring the fluorescence emission intensity as the laser power was modulated. The fluorophores were tested at wavelengths ranging from 750 nm to 1050 nm. The brightest and most stable signals were observed at 1000 nm obtaining values of n = 2.96 *±*0.08 and n = 3.16*±* 0.03 for the blue fluorescing beads and PUREBLU™ Hoechst 33342 dye, respectively. Additionally, their emission spectra were measured and compared to 1P excitation at a laser wavelength of 405 nm (Melles Griot), showing good overlap and corroborating the presence of a 3P signal.

An openSPIM-style, digitally scanned light-sheet fluorescence microscope [16, 17] was implemented for this investigation. The ultrashort pulsed laser beam was expanded to illuminate a single-axis galvanometric mirror (Thorlabs) driven by a triangular wave (Aim-TTi). A virtual light sheet was generated inside the sample chamber by relaying the scanning mirror onto the back aperture of the illumination objective (Nikon, 10x/0.3 numerical aperture (NA), 3.5 mm working distance (wd), water-dipping). Based on measurements of the beam size at the back aperture of the objective, the NA of the light sheet was determined to be 0.17*±*0.01. Samples were held from above and accurately positioned using a XYZ linear translation stage (Newport). Stacks of images were acquired by stepwise motion of the sample across the light sheet using a motorized actuator (PI). Fluorescence was collected perpendicularly to the illumination plane by a second objective lens (Olympus, 20x/0.5 NA, 3.5 mm wd, water-dipping). A 400 mm tube lens (Thorlabs) focused the light on a water-cooled sCMOS camera (ORCA-Flash4.0, HAMAMATSU), yielding a magnification of 40x. Two fluorescence filters (FF01-680/SP, FF01-468/SP, Semrock) were used to block scattered light from the illumination laser and also reject possible undesired 2P signal emitted at longer wavelengths. The microscope can be operated at 2P as well as at 3P by tuning the laser wavelength.

For showing the capability of 3P-LSFM, our first demonstration imaged 1 µm diameter blue fluorescing beads embedded in 1.5 % agarose in a FEP (Fluorinated Ethylene Propylene) capillary. Stacks of images were acquired at steps of 0.25 µm and the performance of the system was compared to 2P. The average laser power was adjusted for each experiment in order to achieve the same maximum intensity values on the camera in both imaging modalities to perform fair comparisons. The laser power available on the sample for the 3P excitation experiments with Gaussian beam illumination at 1000 nm was 259 mW while in the 2P experiments a power of 9.5 mW at a wavelength of 700 nm generated the same fluorescence intensity.

Maximum intensity projections in the axial direction clearly show the intrinsic optical sectioning capability of LSFM (Fig. 1(a, b)). The full width at half maximum (FWHM) of the point spread function (PSF) was measured in various images obtaining an axial resolution of 1.66*±* 0.10 µm and 1.59*±* 0.15 µm for 3P and 2P, respectively (Fig. 1(c)). Approximately, the same axial resolution is achieved in both modalities even using different illumination wavelengths due to the highly confined excitation of the 3P process. The FOV of a light-sheet microscope is usually defined as twice the Rayleigh range of the illumination beam, i.e. the <u>p</u>ropagation range in which the beam width remains less than 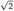 times its minimum size. However, in 3P, the light sheet remains thin *enough* well beyond the expected Rayleigh range due to the properties of the higher order nonlinear excitation process. Consequently, the usable FOV was defined based on the edge-to-edge drop in fluorescence intensity in a 1/e-dependent manner (Fig. 1(d)). Furthermore, the tighter excitation confinement of 3P compared to 2P results in much reduced fluorescence excitation outside the FOV along the propagation direction of the light sheet. For instance, in 3P the usable FOV is 54.6*±* 5.4 µm and the total fluorescence excitation is contained within only 80 µm along the light sheet. In contrast, in 2P the usable FOV is 52.3 *±* 13.7 µm but fluorescence excitation extends up to 140 µm, resulting in additional background fluorescence and photo-damage outside the FOV. It should also be noted that chromatic aberrations in the illumination path make the beam shift both transversally and longitudinally when switching between 3P and 2P. Such strong chromatic aberrations should be accounted for and corrected if simultaneous multicolor experiments are to be performed [18].

**Fig. 1.**
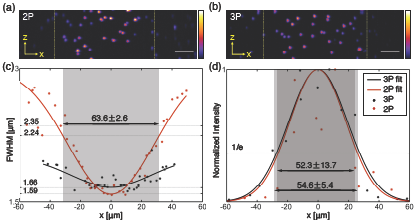
Comparison between 2P- and 3P-LSFM. Axial maximum intensity projections of 3D stacks of images of 1 µm blue fluorescing microspheres embedded in agarose under (a) 2P excitation at 700 nm and (b) 3P excitation at 1000 nm. Scale bar, 10 µm. x-axis: beam propagation; z-axis: optical axis of detection lens. (c) Statistical estimates of the axial resolution and FOV based on FWHM. (d) Statistical estimate of the FOV based on 1/e drop in fluorescence intensity.

The feasibility of using 3P-LSFM for biomedical applications is demonstrated by imaging cellular spheroids of *≈*450 µm in diameter. Human Embryonic Kidney cells (HEK 293 T17) were plated in an ultra-low attachment 96-well round bottom cell culture plate (Corning^®^ Costar^®^ 7007) and grown for 48 hours. After the spheroids were formed, their outer layer was labelled with the PUREBLU™ Hoechst 33342 nuclear staining dye (Fig. 2). Spheroids were embedded in 1% agarose in a FEP capillary. Stacks of images with 0.5 µm spacing were acquired and 3D images were rendered to show the imaging capabilities of the microscope (Supplementary Video). Single 3P slices in the XY and YZ planes are shown in Fig. 2(b, c). In order to assess its performance at depth in scattering samples, the near and far surfaces of the spheroid with respect to the illumination light sheet (blue and red rectangles in Fig. 2(a), respectively) were imaged first for 2P and then for 3P. Stacks were acquired with the same exposure time and the laser power was adjusted to generate equivalent fluorescent signal in the two modalities. Image quality was quantified by measuring the contrast-to-noise ratio (CNR) at various positions in the images [19]. Near the surface, both modalities show the same image quality with similar CNR values as expected (Fig. 2(d, f)). However, at the far surface of the spheroid, 2P-LSFM shows a dramatic drop in image quality (Fig. 2(e)) while 3P-LSFM still preserves high contrast due to the use of a longer wavelength (Fig. 2(g)). The CNR in 2P drops by approximately 71% at a depth of nearly 450 µm while in 3P it only decreases by 15%. Line profiles in Fig. 2(h, i) show the clear improvement in contrast of 3P compared to 2P in imaging at depth (see also **Visualization 1**).

**Fig. 2.**
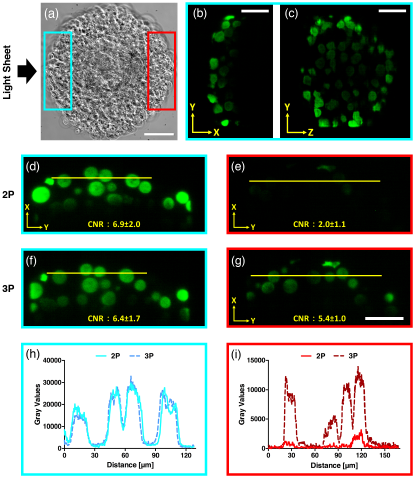
HEK 293 T17 cellular spheroids labeled with PUREBLU™ Hoechst 33342 nuclear staining dye imaged with 2P- and 3P-LSFM. (a) Brightfield image of a spheroid (diameter *≈* 450 µm). The blue and red rectangles represent the near and far surfaces of the spheroid with respect to the light-sheet illumination direction (black arrow). (b, c) Single XY and YZ near-surface planes (blue rectangle in (a)) imaged with 3P-LSFM. (d-g) Single XY planes imaged with 2P-(d, e) and 3P-LSFM in both the near (d, f) and far-surface (e, g). The 3D rendering of the image stacks acquired in both 2P and 3P can be found in the Supplementary Video (see **Visualization 1**). (h, i) Fluorescence intensity profiles along the yellow lines highlighted in (d-g). Average power on the sample for 2P and 3P was 9.5 mW and 259 mW, respectively. Brightfield image scale bar, 100 µm; Fluorescence images scale bar, 50 µm.

To compare our results with theoretical expectations, light-sheet profiles for 2P and 3P were numerically modeled using Fourier beam propagation. In all cases, the following parameters were used: NA= 0.17, *λ*_≈_= 405 nm, *λ*_2*P*_ = 700 nm and *λ*_3*P*_ = 1000 nm. Our simulations of Gaussian light sheets predicted resolutions (given by the FWHM of the light-sheet profile) of 1.5 µm and 1.7 µm and FOV (based on 1/e drop in intensity) of 48 µm and 58 µm for 2P and 3P respectively, which is in agreement with the experimental results (Fig. 1).

Numerical modeling also facilitated exploration of other beam types for 3P-LSFM. Bessel beams have been shown to have much better properties for light-sheet imaging in 2P than 1P [11, 20]. So we also compared Bessel beam illumination for 3P light-sheet microscopy (3P-BB-LSFM). Bessel beams were generated by a thin annulus in the pupil plane of the illumination objective [11]. We define Bessel*β* to denote a Bessel beam generated by an annular ring where *β* is the percentage thickness of the outer radius of the ring. Figure 3(a) shows the cross-sectional light-sheet profiles for a Bessel6.5 beam in 2P and 3P. Due to the extended transverse profile of the Bessel beam, it is not suitable to measure the FWHM to indicate resolution, therefore this was determined from the axial modulation transfer function, *MTF*_*z*_ (*f*_*z*_, *x*) = ℱ_*z*_ (*LS*(*z*, *x*)), where *LS*(*x*, *z*) is the light-sheet cross-section and ℱ_*z*_ denotes the 1D Fourier transform along the axial direction (Fig. 3(b)). The MTF concisely represents information of both resolution and contrast. We set a practical noise-floor at 5% contrast to determine the maximum axial resolution, which is shown in Fig. 3(c). The FOV was determined from the 1/e points in the longitudinal intensity profile of the light sheet (Fig. 3(d)). Figures 3(c,d) show that, for the same NA, 3P-BB-LSFM has a slight reduction in resolution compared to 2P-BB-LSFM but greatly increases the FOV. It also shows that the resolution is effectively decoupled from the FOV as it exhibits very little change with *β*. This can be understood from looking at the cross-section of the light sheet. Figure 3(f) shows the transverse intensity profiles of the light sheets in Fig. 3(a) at ‘focus’ (*x* = 0). For 2P, the contribution of the Bessel beam side-lobes accounts for 24% of the total fluorescence excitation generated on the sample and, when scanned to form the light sheet, these blur into one another giving a broad profile. In 3P the contributions of the side-lobes are suppressed to a greater extent, containing only 4% of the total fluorescence excitation. This makes the light-sheet profile much more Gaussian in shape and so, increasing the propagation-invariant length of the beam (by decreasing *β*) will not significantly affect the resolution. This quantitative MTF study is in agreement with recent works by Chen *et al.* [13] and Rodríguez *et al.* [14] which show benefits of using Bessel beams in 3P confocal microscopy.

**Fig. 3.**
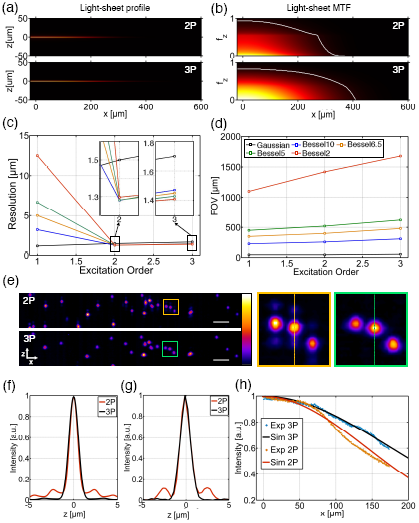
Characterization of LSFM with Bessel beam illumination. (a) Numerically simulated XZ light-sheet cross-sections of a Bessel6.5 beam in 2P and 3P and (b) their respective axial MTFs. The spatial frequency, *f*_*z*_, is normalized to 2*NA*/*λ*_≈_. White lines indicate the isosurface at 5% contrast. (c) Peak axial resolution and (d) FOV for simulated Gaussian and Bessel light-sheets with *β* = 2, 5, 6.5, 10 in 1P, 2P and 3P. Insets in (c) show magnified views of the plot. *λ*_≈_= 405 nm, *λ*_2*P*_ = 700 nm and *λ*_3*P*_ = 1000 nm. (e) Experimental images of 1 µm diameter blue fluorescing beads obtained with 2P- and 3P-BB-LSFM with *β* = 6.5. Average power on the sample for 2P and 3P was 6 mW and 307 mW, respectively. Scale bars, 10 µm. (f) Transverse light-sheet cross-sections at ‘focus’ (*x* = 0) for 2P and 3P obtained from (a). (g) Axial intensity profile of beads in 2P and 3P along the dashed lines in (e). (h) Experimental and simulated longitudinal intensity profile of a Bessel6.5 beam.

A 2P- and 3P-BB-LSFM was implemented experimentally to verify our simulations. A 1° axicon was used to generate an annulus on the back pupil of the illumination objective with *β* = 6.5. Fluorescent beads were imaged (Fig. 3(e)) and their axial intensity profiles are shown in Fig. 3(g). The images obtained in 2P show that the side-lobes are still clearly visible while in 3P their contribution is negligible. Excitation confinement in the main lobe is 80% for 2P and 98% for 3P proving that high aspect ratio light sheets can be generated without the need to use confocal slit detection or deconvolution to eliminate the side-lobes [21]. The intensity profile along the Bessel beam, measured in a Hoechst 33342 dye solution, demonstrates that the use of 3P-BB-LSFM achieves an extended FOV compared to its 2P counterpart (Fig. 3(h)).

Although the average power used in our experiments may be too high for very sensitive biological samples, it can be greatly reduced by using high energy pulses delivered by the above-mentioned optimal laser sources. This, combined with the intrinsic lower photo-damage of LSFM compared to point-scanning microscopy and the reduced photo-damage at longer wavelengths [22], makes 3P-LSFM a promising tool for deep imaging of biological samples.

In summary, we have demonstrated a new LSFM approach based on 3P excitation that results in an extended imaging depth compared with the currently available 2P-LSFM. By imaging *≈*450 µm spheroids, we show that its performance at shallow depths is similar to 2P imaging while at larger depths 3P clearly enables greater image contrast. From our simulations along with the first experimental demonstration of 3P-BB-LSFM, we have shown that the combination of 3P excitation with Bessel beam illumination is even more advantageous for LSFM, achieving deeper penetration and a larger FOV while maintaining high resolution. The penetration depth of the light sheet could be further improved by using longer wavelengths and combining it with attenuation-compensation approaches recently developed for propagation-invariant fields [19]. However, the imaging depth in the axial direction would still be limited by the wide-field detection of visible light which is a continuing avenue of research.

## Acknowledgments

We thank David E. K. Ferrier and Frances Goff for providing samples, Stella Corsetti for assistance with characterization and Roman Spesyvtsev for contributions at early stages of the work.

## REFERENCES

1. W. Denk, J. H. Strickler, and W. W. Webb, Science 248, 73–76 (1990).

2. G. Katona, G. Szalay, P. Maák, A. Kaszás, M. Veress, D. Hillier, B. Chiovini, E. S. Vizi, B. Roska, and B. Rózsa, Nat. Methods 9, 201–208 (2012).

3. N. G. Horton, K. Wang, D. Kobat, C. G. Clark, F. W. Wise, C. B. Schaffer, and C. Xu, Nat. Photonics 7, 205–209 (2013).

4. C. J. Rowlands, D. Park, O. T. Bruns, K. D. Piatkevich, D. Fukumura, R. K. Jain, M. G. Bawendi, E. S. Boyden, and P. T. So, Light. Sci. & Appl. 6, e16255 (2016).

5. D. G. Ouzounov, T. Wang, M. Wang, D. D. Feng, N. G. Horton, J. C. Cruz-Hernández, Y.-T. Cheng, J. Reimer, A. S. Tolias, N. Nishimura, and C. Xu, Nat. Methods 14, 388–390 (2017).

6. F. Helmchen and W. Denk, Nat. Methods 2, 932–940 (2005).

7. J. Huisken, J. Swoger, F. Del Bene, J. Wittbrodt, and E. H. Stelzer, Science 305, 1007–1009 (2004).

8. E. G. Reynaud, U. Kržič, K. Greger, and E. H. K. Stelzer, HFSP J. 2, 266–275 (2008).

9. R. Tomer, K. Khairy, F. Amat, and P. J. Keller, Nat. Methods 9, 755–736 (2012).

10. F. O. Fahrbach, V. Gurchenkov, K. Alessandri, P. Nassoy, and A. Rohrbach, Opt. Express 21, 13824–13839 (2013).

11. T. Vettenburg, H. I. Dalgarno, J. Nylk, C. Coll-Lladó, D. E. Ferrier, T. č ižmár, F. J. Gunn-Moore, and K. Dholakia, Nat. Methods 11, 541–544 (2014).

12. K. Guesmi, L. Abdeladim, S. Tozer, P. Mahou, T. Kumamoto, K. Jurkus, P. Rigaud, K. Loulier, N. Dray, P. Georges, M. Hanna, J. Livet, W. Supatto, E. Beaurepaire, and F. Druon, Light. Sci. Appl. 7, 12 (2018).

13. B. Chen, X. Huang, D. Gou, J. Zeng, G. Chen, M. Pang, Y. Hu, Z. Zhao, Y. Zhang, Z. Zhou, H. Wu, H. Cheng, Z. Zhang, C. Xu, Y. Li, L. Chen, and A. Wang, Biomed. Opt. Express 9, 1992–2000 (2018).

14. C. Rodríguez, Y. Liang, R. Lu, and N. Ji, Opt. Lett. 43, 1914–1917 (2018).

15. C. Xu, W. Zipfel, J. B. Shear, R. M. Williams, and W. W. Webb, Proc. Natl. Acad. Sci. U.S.A. 93, 10763–10768 (1996).

16. P. J. Keller, A. D. Schmidt, J. Wittbrodt, and E. H. Stelzer, Science. 322, 1065–1069 (2008).

17. P. G. Pitrone, J. Schindelin, L. Stuyvenberg, S. Preibisch, M. Weber, K. W. Eliceiri, J. Huisken, and P. Tomancak, Nat. Methods 10, 598–599 (2013).

18. P. Mahou, G. Malkinson, É. Chaudan, T. Gacoin, E. Beaurepaire, and W. Supatto, Small 13, 1701442 (2017).

19. J. Nylk, K. McCluskey, M. A. Preciado, M. Mazilu, Z. Yang, F. J. Gunn-Moore, S. Aggarwal, J. A. Tello, D. E. K. Ferrier, and K. Dholakia, Sci. Adv. 4, eaar4817 (2018).

20. O. E. Olarte, J. Licea-Rodriguez, J. A. Palero, E. J. Gualda, D. Artigas, J. Mayer, J. Swoger, J. Sharpe, I. Rocha-Mendoza, R. Rangel-Rojo and P. Loza-Alvarez, Biomed. Opt. Express 3, 1492–1050 (2012).

21. E. S. Welf, M. K. Driscoll, K. M. Dean, C. Schäfer, J. Chu, M. W. Davidson, M. Z. Lin, G. Danuser and R. Fiolka, Dev. Cell 36, 462–475 (2016).

22. Y. Fu, H. Wang, R. Shi, and J.-X. Cheng, Opt. Express 14, 3942–3951 (2006).

## REFERENCES

1. W. Denk, J. H. Strickler, and W. W. Webb, “Two-photon laser scanning fluorescence microscopy,” Science 248, 73–76 (1990).

2. G. Katona, G. Szalay, P. Maák, A. Kaszás, M. Veress, D. Hillier, B. Chiovini, E. S. Vizi, B. Roska, and B. Rózsa, “Fast two-photon in vivo imaging with three-dimensional random-access scanning in large tissue volumes,” Nat. Methods 9, 201–208 (2012).

3. N. G. Horton, K. Wang, D. Kobat, C. G. Clark, F. W. Wise, C. B. Schaffer, and C. Xu, “In vivo three-photon microscopy of subcortical structures within an intact mouse brain,” Nat. Photonics 7, 205–209 (2013).

4. C. J. Rowlands, D. Park, O. T. Bruns, K. D. Piatkevich, D. Fukumura, R. K. Jain, M. G. Bawendi, E. S. Boyden, and P. T. So, “Wide-field three-photon excitation in biological samples,” Light. Sci. & Appl. 6, e16255 (2016).

5. D. G. Ouzounov, T. Wang, M. Wang, D. D. Feng, N. G. Horton, J. C. Cruz-Hernández, Y.-T. Cheng, J. Reimer, A. S. Tolias, N. Nishimura, and C. Xu, “In vivo three-photon imaging of activity of GCaMP6-labeled neurons deep in intact mouse brain,” Nat. Methods 14, 388–390 (2017).

6. F. Helmchen and W. Denk, “Deep tissue two-photon microscopy,” Nat. Methods 2, 932–940 (2005).

7. J. Huisken, J. Swoger, F. Del Bene, J. Wittbrodt, and E. H. Stelzer, “Optical sectioning deep inside live embryos by selective plane illumination microscopy,” Science 305, 1007–1009 (2004).

8. E. G. Reynaud, U. Kržič, K. Greger, and E. H. K. Stelzer, “Light sheet based fluorescence microscopy: More dimensions, more photons, and less photodamage,” HFSP J. 2, 266–275 (2008).

9. R. Tomer, K. Khairy, F. Amat, and P. J. Keller, “Quantitative high-speed imaging of entire developing embryos with simultaneous multiview light-sheet microscopy,” Nat. Methods 9, 755–763 (2012).

10. F. O. Fahrbach, V. Gurchenkov, K. Alessandri, P. Nassoy, and A. Rohrbach, “Light-sheet microscopy in thick media using scanned bessel beams and two-photon fluorescence excitation,” Opt. Express 21, 13824–13839 (2013).

11. T. Vettenburg, H. I. Dalgarno, J. Nylk, C. Coll-Lladó, D. E. Ferrier, T. č ižmár, F. J. Gunn-Moore, and K. Dholakia, “Light-sheet microscopy using an airy beam,” Nat. Methods 11, 541–544 (2014).

12. K. Guesmi, L. Abdeladim, S. Tozer, P. Mahou, T. Kumamoto, K. Jurkus, P. Rigaud, K. Loulier, N. Dray, P. Georges, M. Hanna, J. Livet, W. Supatto, E. Beaurepaire, and F. Druon, “Dual-color deep-tissue three-photon microscopy with a multiband infrared laser,” Light. Sci. Appl. 7, 12 (2018).

13. B. Chen, X. Huang, D. Gou, J. Zeng, G. Chen, M. Pang, Y. Hu, Z. Zhao, Y. Zhang, Z. Zhou, H. Wu, H. Cheng, Z. Zhang, C. Xu, Y. Li, L. Chen, and A. Wang, “Rapid volumetric imaging with Bessel-Beam three-photon microscopy,” Biomed. Opt. Express 9, 1992–2000 (2018).

14. C. Rodríguez, Y. Liang, R. Lu, and N. Ji, “Three-photon fluorescence microscopy with an axially elongated Bessel focus,” Opt. Lett. 43, 1914–1917 (2018).

15. C. Xu, W. Zipfel, J. B. Shear, R. M. Williams, and W. W. Webb, “Multiphoton fluorescence excitation: new spectral windows for biological nonlinear microscopy,” Proc. Natl. Acad. Sci. U.S.A. 93, 10763–10768 (1996).

16. P. J. Keller, A. D. Schmidt, J. Wittbrodt, and E. H. Stelzer, “Reconstruction of Zebrafish Early Embryonic Development by Scanned Light Sheet Microscopy,” Science. 322, 1065–1069 (2008).

17. P. G. Pitrone, J. Schindelin, L. Stuyvenberg, S. Preibisch, M. Weber, K. W. Eliceiri, J. Huisken, and P. Tomancak, “OpenSPIM: an open-access light-sheet microscopy platform,” Nat. Methods 10, 598–599 (2013).

18. P. Mahou, G. Malkinson, É. Chaudan, T. Gacoin, E. Beaurepaire, and W. Supatto, “Metrology of Multiphoton Microscopes Using Second Harmonic Generation Nanoprobes,” Small 13, 1701442 (2017).

19. J. Nylk, K. McCluskey, M. A. Preciado, M. Mazilu, Z. Yang, F. J. Gunn-Moore, S. Aggarwal, J. A. Tello, D. E. K. Ferrier, and K. Dholakia, “Light-sheet microscopy with attenuation-compensated propagation-invariant beams,” Sci. Adv. 4, eaar4817 (2018).

20. O. E. Olarte, J. Licea-Rodriguez, J. A. Palero, E. J. Gualda, D. Artigas, J. Mayer, J. Swoger, J. Sharpe, I. Rocha-Mendoza, R. Rangel-Rojo and P. Loza-Alvarez, “Image formation by linear and nonlinear digital scanned light-sheet fluorescence microscopy with gaussian and bessel beam profiles,” Biomed. Opt. Express 3, 1492–1505 (2012).

21. E. S. Welf, M. K. Driscoll, K. M. Dean, C. Schäfer, J. Chu, M. W. Davidson, M. Z. Lin, G. Danuser and R. Fiolka, “Quantitative Multiscale Cell Imaging in Controlled 3D Microenvironments.” Dev. Cell 36, 462–475 (2016).

22. Y. Fu, H. Wang, R. Shi, and J.-X. Cheng, “Characterization of photo-damage in coherent anti-Stokes Raman scattering microscopy,” Opt. Express 14, 3942–3951 (2006).

